# Integrative multi-omics reveals PARP14 as a key IFNγ-regulated mediator of metastatic progression in Ewing sarcoma

**DOI:** 10.64898/2025.12.23.696151

**Authors:** Sara Sánchez-Serra, Mariona Chicón-Bosch, Núria Martorell-Villanueva, Eduardo Candeal, Ignasi Jarne-Sanz, Joan Josep Bech-Serra, Daniel Giménez-Llorente, Manuel Gris-Lorente, Paola Monaco, Laura Santana-Viera, Silvia Mateo-Lozano, Jaume Mora, Mario Niepel, Carolina de la Torre Gomez, Rebeca Sanz-Pamplona, Ana Cuadrado, Ana Losada, Roser López-Alemany, Òscar M Tirado

## Abstract

Metastatic Ewing sarcoma (EwS) remains a major clinical challenge, highlighting the need to identify molecular drivers of disease dissemination. Through integrated transcriptomic, functional, and mechanistic analyses, we identify PARP14 as a key regulator of EwS metastatic progression. PARP14 is consistently upregulated in metastatic lesions across spontaneous and xenograft mouse models as well as in patient samples. Genetic and pharmacologic depletion of PARP14 (via siRNA knockdown, CRISPR-Cas9 knockout, or PROTAC-mediated degradation) significantly impaired EwS cell migration and invasion in vitro, without affecting proliferation or clonogenic capacity. In vivo, PARP14 loss delayed tumor growth and reduced metastatic burden, supporting a functional role in disease progression. Conversely, IFNγ-induced PARP14 upregulation and stable overexpression enhanced EwS cell migration and increased lung metastasis formation.

Transcriptomic profiling of PARP14-deficient cells revealed downregulation of invasive signatures and enrichment of inflammatory and p53-related pathways, together with gene sets associated with chromatin remodeling. Subcellular fractionation and interactome analyses showed that PARP14 localizes to cytoplasmic, nuclear, and chromatin-associated compartments, and interacts with chromatin, cytoskeletal and ECM regulators. ChlP analyses further demonstrated PARP14 binding to promoters enriched for NRF1 motifs, suggesting a role as a transcriptional co-regulator influencing gene networks relevant to EwS progression.

Together, our results establish PARP14 as a contributor to EwS dissemination and uncover a previously unappreciated chromatin-associated role for this protein, supporting its potential as both a biomarker and therapeutic target in metastatic EwS.

**STATEMENT OF SIGNIFICANCE:** We identify PARP14 as a novel driver of Ewing sarcoma metastasis through integrated multi-omics profiling. Genetic and pharmacological interrogation establishes PARP14 as a promising therapeutic target for this understudied malignancy.

## INTRODUCTION

Ewing sarcoma (EwS) is the second most common primary bone malignancy in children and young adults^1^. It is a highly aggressive tumor associated with poor clinical outcomes, particularly for metastatic patients, whose 5-year survival rate remains below 30%^2^. Despite intensive multimodal treatment including chemotherapy, surgery and radiotherapy, the prognosis for patients with metastatic or relapsed EwS remains dismal^3^. A key obstacle to improving outcomes is the limited understanding of the molecular mechanisms driving metastasis. Identifying the critical regulators of metastatic progression is therefore essential for the development of more effective therapeutic strategies to improve patient survival and quality of life.

Previous transcriptomic analysis comparing paired primary tumors and metastases from a spontaneous metastasis EwS mouse model identified PARP14 as significantly upregulated in metastatic lesions^4^. PARP14 is the largest member of the poly(ADP-ribose) polymerase (PARP) family, enzymes that catalyze ADP-ribosylation of their targets thereby regulating a wide range of cellular processes, including DNA repair, transcription, and signal transduction^5^. Interest in PARP14 has grown in recent years as a potential drug target for inflammation and cancer, given its regulatory roles in immune signaling and tumor progression^6^. Indeed, several small-molecule inhibitors targeting this protein have been purposed as a therapeutic strategy^7^. Notably, PARP14 has been implicated in shaping the tumor microenvironment (TME) by promoting pro-tumor macrophage polarization as well as suppressing antitumor inflammation response by modulating IFN-y and IL-4 signaling^8–10^. Additionally, PARP14 contributes to chemotherapy and immune checkpoint blockade, and its inhibition has been shown to sensitize certain tumors to these therapies^11^.

Here, we demonstrate that PARP14 is consistently upregulated in EwS metastases and that its elevated expression correlates with enhanced migration and invasion of EwS cells both *in vitro* and *in vivo*. Remarkably, these pro-metastatic effects appear to occur independently of PARP14’s catalytic activity. We also describe its localization to the nucleus, where it interacts with the histone methyltransferase SETDB1 and binds to the promoter regions of genes regulated by NRF1, suggesting a role as a transcriptional co-regulator influencing gene networks relevant to EwS progression. Altogether, our findings identify PARP14 as a key regulator of EwS metastatic progression and highlight its potential as both a biomarker and a therapeutic target in this aggressive pediatric malignancy.

## MATERIALS AND METHODS

### Cell lines and culture conditions

EwS cell lines A673 and TC252 were gifted from Dr. Heinrich Kovar (St. Anna Children’s Cancer Research Institute, Vienna, Austria); MHHES1 was purchased from the German Collection of Microorganisms and Cell Cultures (DSMZ, Braunschweig, Germany). Authenticity of the cell lines was routinely confirmed by STR profiling analysis.

Cell lines were cultured in RPMI 1640-GlutaMAX medium (Gibco, Thermo Fisher Scientific, Carlsbad, CA, USA) supplemented with 10% (v/v) fetal bovine serum (FBS; Gibco) and 1% (v/v) penicillin-streptomycin (P/S; Gibco). Cells were incubated at 37 °C in a humidified atmosphere of 5% CO_2_ in air and checked regularly for mycoplasma contamination using DNA staining with Hoechst 33342 dye (Thermo Fisher Scientific) or by agarose gel electrophoresis of PCR products. Exponentially growing cells were used for all experiments.

A673 transfected with luciferase expressing construct (A673Luc; gift from Dr. lbane Abasolo, VHIR, Barcelona), was cultured in antibiotic selection media (puromycin; Sigma-Aldrich, Saint Louis, MO, USA). These cells were used for the *in vivo* experiments as they could be detected and imaged using luciferase activity through *In Vivo* Imaging System (IVIS®; PerkinElmer, Shelton, CT, USA).

Metastasis-derived primary cultures from A673Luc and TC252 mice lung metastasis were obtained (see below). Lung metastatic tissue was placed in cell culture plates, ground with a scalpel and incubated for 2 h at 37 °C on a shaking incubator in RPMI + collagenase II 0.1% (Gibco). Same volume of RPMI + 20% FBS + 1% P/S was added and filtered using a 70 µm Nylon filter (BO Falcon, Schaffhausen, Switzerland) and any tissue remaining on the filter was scraped off. Tissue was centrifuged at 3500 g for 10 min, resuspended in RPMI + 20% FBS + 1% P/S and placed in a cell culture plate at a confluency of 1-2 × 10^6^ cells/p60 plate. Once cells reached 80% confluency, they were cultured as standard cell lines (described before). Primary cultures obtained from A673Luc lung metastases named as Met1-2-3A6; those obtained from TC252 lung metastases named as Met2-3-4-5TC.

### Spontaneous metastasis mouse model

Orthotopic model for metastasis detection in mice was performed as described12 using 6-week-old female, athymic nude mice Hsd:Athymic *Mice-Foxn1*^*nu*^ from Envigo (Indianapolis, IN, USA). Briefly, luciferase-labeled A673 EwS cells (A673Luc; 1 × 10^6^ cells) or TC252 EwS cells (not luciferase-labeled; 2.5 × 10^6^ cells) resuspended in 100 μL PBS (Sigma-Aldrich) were injected into the gastrocnemius muscle of mice. Tumor size was monitored using calipers. Once the tumor reached approximately 800 mm^3^, around day 15 –20, the muscles containing the tumor were surgically resected (evaluation by size/weight and also IVIS lecture for A673Luc). After surgery, mice could survive for a period long enough to enable the development of distant metastases. The presence of morbidity symptoms was checked every 48 h. At day 90 after cell inoculation, lung metastases were detected by *in vivo* lecture of luminescence (IVIS) of the whole mouse (for A673Luc) or *ex vivo* once lungs were resected. Mice were euthanized, and lungs were extracted, fixed in 4% paraformaldehyde and embedded in paraffin. Lung sections were stained with hematoxylin and eosin and metastases were counted under an optical microscope (Nikon i80). Some of the mice developed local relapses in areas adjacent to the surgery site and were euthanized before the experiment end point but also included in the analyses.

Animals were cared according to the Institutional Guidelines for the Care and Use of Laboratory Animals. All mice were kept under Specific Pathogen Free conditions in standard polycarbonate cages, 5 mice/cage, and fed sterile food and water available *ad libitum*. Facilities were maintained at a controlled temperature (21-23 °C) and humidity (50–60%) and kept on a 12/12 h light/dark cycle. Ethics approval was provided by a locally appointed ethics committee from IDIBELL (Barcelona, Spain). IDIBELL animal facility abides by the Association for Assessment and Accreditation of Laboratory Animal Care (AAALAC) regulations.

### Transcriptomic array and analysis (Clariom™ D array)

Affymetrix Clariom™ D transcriptomic array was performed as previously described^4^. Briefly, total RNA (2 µg) was extracted from samples (n = 20) using the NucleoSpin RNA II (mRNA) from Macherey-Nagel (Düren, Germany), following manufacturer’s instructions including a DNase treatment step. Quantification and first quality control were performed on a NanoDrop ND-1000 (Thermo Fisher Scientific, Waltham, MA, USA). Quality control assessment with Agilent 2100 Bioanalyzer (Agilent Technologies, Santa Clara, CA, USA) was performed before microarray analysis. Only samples with RNA Integrity Number (RIN) above 7 were processed. Gene expression microarray was performed using Clariom D Array (Thermo Fisher Scientific) at Servei d’Anà lisi de Microarrays (MARGenomics, lnstitut Hospital del Mar d’lnvestigacions Mediques (IMIM), Barcelona, Spain). Amplification, labeling and hybridization were performed according to the GeneChip WT PLUS Reagent kit and the samples were hybridized to Clariom D Human (Thermo Fisher Scientific) in a GeneChip Hybridization Oven 640. Washing and scanning were performed using the Expression Wash, Stain, and Scan Kit (Affymetrix Inc., Thermo Fisher Scientific).

Gene expression (Clariom D) array data was analyzed at the Sarcoma Research Group. After quality control of raw data (aroma.affymetrix package^13^), samples were background corrected and normalized using the robust multi-chip average (oligo package^14^) method. In order to detect differentially expressed genes (DEGs) between sample groups, the Linear Models for Microarray (limma^15^) package was used. Correction for multiple comparisons was performed using false discovery rate (FDR). Transcripts with adjusted p-value <0.05 (or p-value <0.05 when stated) and absolute fold-change (FC) above 1.5 (absolute log2FC (LFC) > 0.58) were selected as significant. Figures were obtained using ggplot2 package^16^ and RCOLORBREWER package^17^. Analyses were performed in R (v3.5.2, http://www.R-project.org/).

### Reverse transcription quantitative PCR (RT-qPCR)

Total RNA was extracted using the NucleoSpin RNA II (mRNA) kit (Macherey-Nagel) following manufacturer’s instructions including a DNase treatment step. After mRNA quantification using NanoDrop ND-1000, 2 µg RNA was used for complementary DNA synthesis with M-MLV Reverse Transcriptase (lnvitrogen, Thermo Fisher Scientific). Quantitative reverse transcription-PCR (qRT-PCR) was performed under universal cycling conditions on LightCycler 480 II (Roche, Basel, Switzerland) using TaqMan PCR Mastermix and TaqMan probes (Applied biosystems, Thermo Fisher Scientific; PARP14 Hs01395601_m1, PPIA Hs04194521_m1). Cycle threshold (CT) values were normalized to that of PPIA. Relative expression levels of the gene of interest were calculated using the ΔΔCT (M.W. Pfaffl^18^) mathematical model.

### Protein extraction and western blot

Protein analysis by western blot (WB) was performed as previously described^19^. Primary antibodies used were PARP14 1:400 #sc-377150 (Santa Cruz Biotechnology, Dallas, TX, USA). Secondary antibodies used were horseradish peroxidase-conjugated goat anti-rabbit #P0448 and goat anti-mouse #P0260 (Dako, Agilent Technologies, Santa Clara, CA, USA). Peroxidase activity on membranes was detected by enhanced chemiluminescence (Pierce, Thermo Fisher Scientific) following manufacturer’s instructions. Tubulin 1:10000 #6199 (Sigma-Aldrich) was used as loading control.

### Clinical material

All patient samples were collected following written informed consent and approval by the Institutional Ethics Review Committee. Fresh tumor specimens were obtained from surgical resections at Hospital Sant Joan de Deu (HSJD; Esplugues de Llobregat, Barcelona, Spain). Ewing sarcoma diagnosis was confirmed by board-certified pathologists and molecular biologists using standardized assays, including RT-PCR and FISH for fusion gene detection when applicable.

### Immunohistochemistry (IHC)

After dewaxing, expression of PARP14 in mice xenografts and patient samples was analyzed using a rabbit polyclonal antibody (#HPA008846, Sigma-Aldrich). Sections were then dehydrated and mounted using DPX non-aqueous mounting medium (Sigma-Aldrich). Images were taken with a Nikon Eclipse 80i microscope.

### Transient gene silencing

For transient gene silencing, cells were transfected with DharmaFECT (Dharmacon, Lafayette, CO, USA) following manufacturer’s protocol and using 100 nM of a mixture of 2 customized small interfering RNAs (siRNAs) against PARP14’s sequence (siPARP14_1 5’-AGGCCGACUGUGACCAGAUAGUGAA-3’ and siPARP14_2 5’ CGGCACUACACAGUGAACUUGAACA-3’). As negative control, a customized non-targeting siRNA was used siNT 5’-UAAGGCUAUGAGAGAUAC-3’).

### Stable transfections

To achieve stable PARP14 silencing, the non-homology mediated (KN2.0) CRISPR knock-out kit (#KN420878 Origene, Rockville, MD, USA) was used. Cells were transfected with Lipofectamine 2000 (#11668030 Life Technologies, Thermo Fisher Scientific) following manufacturer’s protocol and using 1 µg of guide RNA (gRNA) or scrambled control vector (SCR) and 1 µg of donor DNA.

Stable gene overexpression was obtained by transfection using Lipofectamine 2000 following manufacturer’s protocol and using 1 µg of PARP14-FLAG-His vector containing a BFP2 reporter (VectorBuilder customed design) or CMV control vector (VectorBuilder).

For clonal selection, pools were seeded on a 96-well plate at a confluence of 1 cell per well in complete medium with the correspondent selection antibiotic (puromycin (Sigma-Aldrich) / neomycin (Life Technologies, Thermo Fisher Scientific)). Cells were allowed to grow for 2 weeks and resistant clones were isolated and tested for RNA and protein expression analysis to confirm PARP14 expression.

### Cell treatments

For PARP14 protein degradation, proteolysis targeting chimeras (PROTACs; gifted by RISON Therapeutics, Cambridge, MA, USA) were used. The day after seeding, Compound 3 (RBN012811; patent #CA3142002A1) its enhanced and more selective version Compound A or the inactive PROTAC Compound 2 (RBN013527) were added to the cells at 10 nM in complete medium. DMSO was used as a vehicle control. Cells were collected after 24-48 h depending on downstream experiment (protein expression or phenotypic validation).

For PARP14 catalytic function inhibition, the enzymatic inhibitor Compound 8 (RBN012759) or its inactive form Compound 1 (gifted by RISON Therapeutics) were used. The day after seeding, the compounds were added to the cells at 1 µM in complete medium. DMSO was used as a vehicle control. Cells were collected after 24-48 h depending on downstream experiment.

For IFNγ treatments, complete medium containing 100 ng/ml of recombinant human IFNγ (PeproTech) was added the day after seeding. Phosphate buffered saline (PBS; Biowest, Nuaillé, France) containing 0.1% Bovine serum albumin (BSA, Thermo Fisher Scientific) was used as a vehicle control. Cells were collected after 3-24 h depending on downstream experiment.

### Transcriptomic array and analysis (RNA sequencing)

Total RNA (2 µg) was extracted from samples (n = 12) using the NucleoSpin RNA II (mRNA) from Macherey-Nagel following manufacturer’s instructions including a DNase treatment step. Quantification and first quality control were performed on a NanoDrop ND-1000 (Thermo Fisher Scientific). Truseq stranded mRNA Library + Novaseq 6000 150PE (150×2 bp) 20M total reads (3 Gb/spl) was performed at Macrogen (Seoul, South Korea). 1 µg of RNA for each sample in triplicate was sent to Macrogen, where the quality control and quantification were tested again before sequencing.

Bioinformatic analysis of the obtained results was performed in collaboration with the Institute de lnvestigación Sanitaria Aragon (IIS Aragón; Zaragoza, Spain). RNAseq raw data were processed on an external high-performance computing server. Raw reads were assessed with FastQC v0.12.1 and trimmed using BBduk (BBMap v39.01). Clean reads were aligned to the human genome GRCh38.p13 with GENCODE v40 annotation using STAR v2.7.10b. Gene and transcript quantification was performed with RSEM v1.2.28, and quality metrics were summarized with MultiQC v1.32. Count data were imported into R v4.3.0 for differential expression analysis using DESeq2 v1.42.1. Statistical significance was defined as p-value ≤ 0.005 and absolute LFC ≥ 1, and differentially expressed genes were ranked accordingly.

Visualization of results, including heatmaps and exploratory plots, were generated using pheatmap, ggplot^16^ and RcolorBrewer packages^17^. All analyses were performed in R (v4.3.0, http://www.R-project.org/) unless otherwise specified.

Functional analysis was performed on the Gene Set Enrichment Analysis (GSEA) desktop software using the pre-ranked tool^20^, using collections **H** (Hallmark) and C2 (Curated gene set; KEGG).

### Boyden chamber assays

For Transwell migration assay, cells were harvested and centrifuged for 5 min at 1500 g. After an additional wash with RPMI, 1.5 x 105 cells in 150 **µL** serum-free medium was added to the top chamber of 8-µm pore polycarbonate transwells (Transwell Permeable Supports, #353097 Falcon). In the bottom chamber, 500 µL complete medium (10% FBS) was added and used as a chemoattractant. After 24-48 h (cell line dependent) cells on the upper chamber were removed with a cotton swab. Migrated cells on the membrane underside were fixed for 30 min using 70% ethanol and stained with 2% crystal violet for 20 min. Transwell membranes were collected and 4 pictures of each membrane were acquired by optical microscopy (10×). Migrated cells were counted manually using lmageJ software^21^. Alternatively, cells were discolored with a 10% glacial acetic acid solution and crystal violet was quantified by spectrometry (*λ* = 570 nm).

For Transwell invasion assay, the same protocol was followed but transwells were previously coated with 50 µL cold Matrigel (BD Biosciences, Franklin Lakes, NJ, USA) diluted 1:20 in RPMI and placed in a 37 °C incubator for 6 h. After Matrigel polymerization, cells were seeded, fixed, stained, and counted as described. For this assay, 2 × 10^5^ cells in 200 µL serum-free medium were seeded to the transwells.

### Cell proliferation assays (MTT)

For viability assays, 1000 cells were seeded in 96-well plates. At 24, 48, and 72 h after seeding, culture medium was removed and 100 µL of water-soluble tetrazolium (WST-1, Roche) diluted 1:20 in complete medium was added to each well. After 3 h of incubation at 37 °C, cell viability was quantified by spectrometry (λ = 440 nm).

### Colony formation assays

For clonogenic assays, 500-1000 cells (cell line dependent) were seeded in 6-well plates. When colonies reached saturation, approximately 14 days after seeding, cells were fixed with cold methanol for 10 min, washed with PBS (Biowest), stained with 2% crystal violet (Sigma-Aldrich) for 20 min, and washed with water. Plates were then scanned and colonies were counted manually using Image J software. Alternatively, colonies were discolored with a 10% glacial acetic acid solution and crystal violet was quantified by spectrometry (λ = 570 nm).

### Sub-cellular fractionation

Chromatin fractionation was performed as described^22^. Cells were resuspended at 2 × 10^7^ cells/ml in buffer A (10 mM HEPES pH 7.9, 10 mM KCI, 1.5 mM MgCl2, 0.34 M sucrose, 10% glycerol, 1 mM OTT, 1 mM NaV04, 0.5 mM NaF, 5 mM β-glycerophosphate, 0.1 mM PMSF), and incubated on ice for 5 min in the presence of 0.05% Triton X-100. Low-speed centrifugation (4 min/600 g/4°C) allowed the separation of the cytosolic fraction (supernatant) and nuclei (pellet). Nuclei were washed and subjected to hypotonic lysis in buffer B (3 mM EOTA, 0.2 mM EGTA, 1 mM OTT, 1 mM NaV04, 0.5 mM NaF, 5 mM β-glycerophosphate, 0.1 mM PMSF) for 30 min on ice. Nucleoplasmic and chromatin fractions were separated after centrifugation (4 min/600 g/4°C). Chromatin was resuspended in Laemmli buffer and sonicated twice for 15 seconds at 20% amplitude and prepared for immunoblotting.

### Immunofluorescence (IF)

To analyze PARP14 location by immunofluorescence, cells were seeded in 4-well glass chamber slides (#PEZGS0416, Millipore, Sigma-Aldrich). Transfected PARP14 Kl cells contained a BFP2 reporter. Nuclei were stained using BioTracker 650 Red Nuclear dye, (#SCT119, Sigma-Aldrich). Images were taken with a confocal Leica TCS SP5 microscope (Leica Microsystems, Wetzlar, Germany).

### Pull-down

Cells expressing PARP14_FLAG-His tagged vector or an empty CMV FLAG tagged vector were harvested per standard protocol (from 150 mm culture dishes). Cell pellets were lysed with 500 µL lysis buffer (250 mM KCI, 1 mM EOTA, 1% Triton, 0.05% NP-40, 1 mM OTT, 10% glycerol, 50 mM Tris pH 7.8 supplemented with protease and phosphatase inhibitors) and incubated with Benzonase nuclease (75 activity units/sample; Sigma-Aldrich) for 6 h at 4 °C with agitation. Subsequently, samples were centrifuged for 10 min at 16200 g 4 °C.

Immunocomplexes were purified using the anti-FLAG M2 affinity gel (Sigma-Aldrich). Prior to incubation, 40 µL per sample of anti-FLAG beads were washed thrice in BC100 buffer (100 mM KCI, 0.05 mM EOTA, 0.05% NP-40, 10% glycerol, 1 mM OTT and 10 mM Tris; pH 7.8). Protein extracts were then incubated with anti-FLAG beads overnight at 4 °C with agitation. The following day, beads were washed thrice in BC500 buffer (500 mM NaCl, 0.05 mM EOTA, 0.05% NP-40, 10% glycerol, 1 mM OTT and 10 mM Tris; pH 7.8). Bound proteins were eluted based on competent binding with 200 ng/ml FLAG peptide (Sigma-Aldrich). Eluates were mixed with loading buffer, denatured 10 min at 100 °C and analyzed by WB.

### Mass spectrometry

After pull down, agarose beads were sent dry after BC500 washes to the Proteomics Unit at Josep Carreras Leukaemia Research institute (Badalona, Barcelona, Spain) for its mass spectrometry (MS) analysis. The beads were then washed 3 times with PBS 1X (500 µL of PBS + centrifuge 1000 g for 5 min) and 3 times with Tris 0.1 M (same procedure as PBS). After discarding the supernatant (Tris 0.1 M), the beads were resuspended with 6 µL of Urea 6 M / Tris 0.1 M. Reduction was performed with OTT 10 mM in Tris 0.1 M for 1 h at 30 °C and shaking 650 rpm. Alkylation was performed in the dark with 2-chloroacetamide 55 mM in Tris 0.1 M for 30 min, at 30 °C, shaking 650 rpm. Samples were then diluted with 280 µL of Tris 0.1 M and 5 µL of trypsin 0.2 µg/µL were added to digest the proteins overnight at 30°C and shaking 650 rpm. Finally, samples were centrifuged for 5 min and 5000 g. The beads were discarded. The digestion was stopped with 20 uL Formic Acid 100%, dried in a speed-vac and stored at −80 °C until use.

Dried-down peptide samples were reconstituted with 0.1% FA and analyzed by reverse-phase LC-MS/MS using an Evosep One LC coupled to an Exploris 480 mass spectrometer (Thermo Fisher Scientific) by direct data-independent (dDIA) approach. Peptides were resolved using the SPD30 method (44 min) using an E1106 performance column (15 cm, 150 µm, 1.9 µm beads; Evosep). Sample data was acquired in a data-independent mode (DIA) with a full MS scan (scan range: 350 to 1000 m/z; resolution: 120000; maximum injection time: 50 ms; normalized AGC target: 300%) and 50 periodical MS/MS segments, applying 10 Th isolation windows (0.5 Th overlap; resolution: 30000; maximum injection time: 50 ms; normalized AGC target: 1000%). Peptides were fragmented using a normalized HCD collision energy of 30%, acquiring data in profile and centroid mode for full MS and MS/MS scans, respectively.

RAW Thermo files were analyzed using DIA-NN version 1.8.1. A *Homo sapiens* protein database downloaded from UniProt on February 6, 2024, containing isoforms and restricted to curated entries (20397 entries), was used in DIA-NN to predict the spectral library for peptide identification. Methionine oxidation and protein N-terminal acetylation were set as variable modifications, while carbamidomethylation of cysteine residues was specified as a fixed modification. A maximum of two missed cleavages and one variable modification per peptide were allowed. The output was filtered at a false discovery rate (FDR) of 0.01 at PSMs, peptides and protein groups level.

For interactome analysis, SAINTq software (version 0.0.4) was used with default parameters. The DIA-NN protein group output file (report.pg_matrix.tsv) was reformatted according to the software specifications and used to identify potential interacting candidates.

### Chromatin immunoprecipitation (ChlP)

A673 cells were crosslinked with 1% formaldehyde for 15 min at room temperature. After quenching the reaction with 0.125 M Glycine, fixed cells were washed twice with PBS containing 1 µM PMSF and protease inhibitors. Cells were then lysed in lysis buffer (1% SDS, 10 mM EDTA, 50 mM Tris-HCI pH 8.1) at a concentration of 2×10^7^ cells/ml. Sonication was performed using a Covaris E220 evolution (shearing time 7 min at 5-7 °C range with 140 peak incident power, 5% duty factor and 200 cycles per burst) in a volume of 1 ml. Chromatin from 10^7^ cells was incubated with 20 µg of the antibody as described^23^. For library preparation, at least 5 ng of DNA were processed through subsequent enzymatic treatments using “NEBNext Ultra II FS DNA Library Prep Kit for lllumina”. Briefly, a short fragmentation of 10 min was followed by end-repair, dA-tailing, and ligation to adapters. The adapter-ligated libraries were completed using limited-cycle PCR (10 cycles). The resulting average fragment size was 300 bp, of which 120 bp corresponded to the adapter sequences. Libraries were applied to an lllumina flow cell for cluster generation and sequenced on an lllumina NextSeq 500 (with v2.5 reagent kits) following manufacturer’s recommendations.

### Statistical analysis

Data was analyzed for statistical significance using Student’s t-test or analysis of variance with Bonferroni’s correction. Fisher’s exact test was used for evaluating differences in lung metastasis incidence in mice. Data represented in Kaplan-Meier plots was analyzed with Mantel-Cox test. Threshold for significance was p-value <0.05. Statistical analyses were performed using GRAPHPAD PRISM v9.4.0 (GraphPad software, Sant Diego, CA, USA). Unless otherwise stated, experiments were performed thrice. Statistical differences in graphs as: *p ≤ 0.05, **p ≤ 0.01, ***p ≤ 0.001, ****p ≤ 0.0001.

## RESULTS

### PARP14 is overexpressed in EwS metastases

To gain insight into the molecular mechanisms underlying EwS metastatic progression, transcriptomic analysis (Clariom D^™^ Array) was performed as described before, comparing primary tumors and metastases from our spontaneous metastasis mouse model. Among the differentially expressed genes (DEGs), PARP14 emerged as one of the most significantly upregulated in metastatic lesions (Log2 fold-change (LFC) = 2.18; Fig. S1A).

To validate these findings, we analyzed the same cohort used for the transcriptomic profiling along with an independent cohort that included additional tumor and metastasis samples. In 6 out of 7 matched pairs (85%), PARP14 expression was significantly higher in metastases relative to their corresponding primary tumors (Fig. 1A). To ensure this was not a cell line-dependent phenomenon, we also assessed PARP14 expression in primary tumors and metastases derived from TC252 xenografts, confirming consistent upregulation in metastatic samples in this independent model (Fig. S1B).

**Figure 1.**
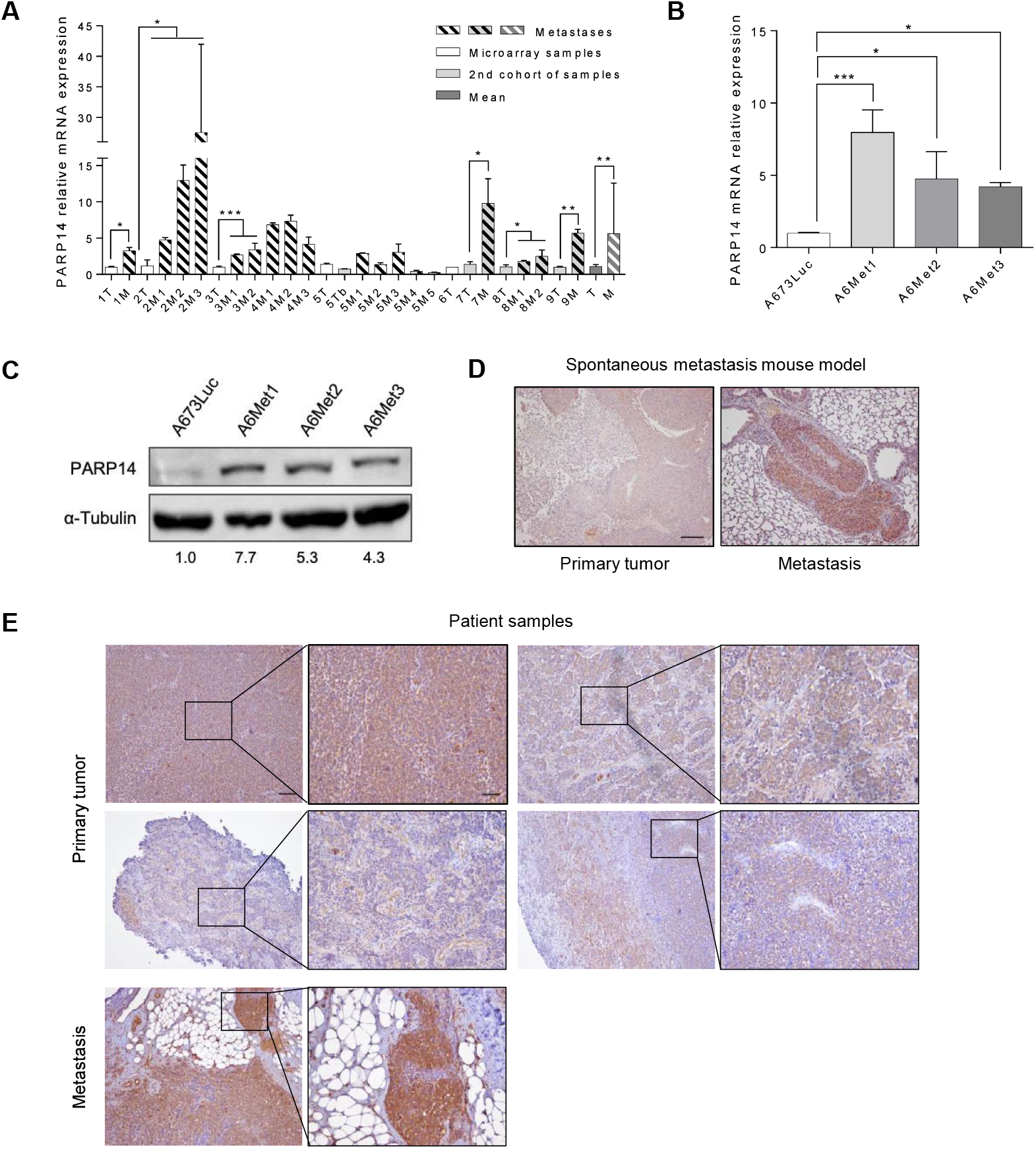
PARP14 is overexpressed in EwS metastases. **A)** PARP14 mRNA relative levels of both microarray and extra EwS xenograft samples by RT-qPCR. mRNA levels of each metastasis are relative to their corresponding tumor levels (normalized to 1). T: tumor; M: metastasis. Means of both group samples are also shown. **B)** PARP14 Mrna relative levels on A673 xenograft-derived metastatic cell lines (A6Met1-3) by RT-qPCR. mRNA levels of each metastatic cell line are relative to A673Luc (cells injected to the metastatic mouse model, normalized to 1). C) PARP14 protein relative levels on A673 xenograft-derived metastatic cell lines by WB. α-Tubulin was used as loading control. PARP14 protein levels of each metastatic cell line are relative to A673Luc. **D)** PARP14 protein expression assessed by immunohistochemistry staining of primary tumor and lung metastasis from xenograft mouse model. Scale bar= 50 μm. **E)** PARP14 protein expression assessed by immunohistochemistry staining of primary tumors and a metastatic sample from a small cohort of EwS patients. Scale bar = 50 μm. Statistical analyses were made using unpaired t-test with Welch’s correction and one-way ANOVA test. ^*^p<0.05; ^**^p<0.01; ^***^p<0.001.

We further evaluated PARP14 expression in threeA673 xenograft-derived metastatic cell lines (A6Met1– 3), established as primary cultures from mouse lung metastases. Consistently, these metastatic cell lines exhibited significantly elevated PARP14 expression compared with the parental A673Luc cell line used for engraftment (Fig. 1B–C). In addition, we confirmed PARP14 upregulation *in situ* in mouse metastatic tissues (Fig. 1D) and in a small cohort of EwS patient samples (Fig 1E).

### Loss of PARP14 impairs migration and invasion in EwS

To investigate the functional role of PARP14 in EwS metastasis, loss-of-function transwell migration assays were performed. Transient PARP14 knock-down using siRNAs (Fig. 2A; Fig. S2A) significantly reduced migration ability of two cell lines derived from patient metastases, TC252 and MHHES1 (Fig. 2B; Fig. S2B). Consistently, CRISPR-Cas9 mediated PARP14 knock-out (KO, Fig. 2C; Fig. S2C) also translated to an impaired migration ability in these cells (Fig. 2D; Fig. S2D).

**Figure 2.**
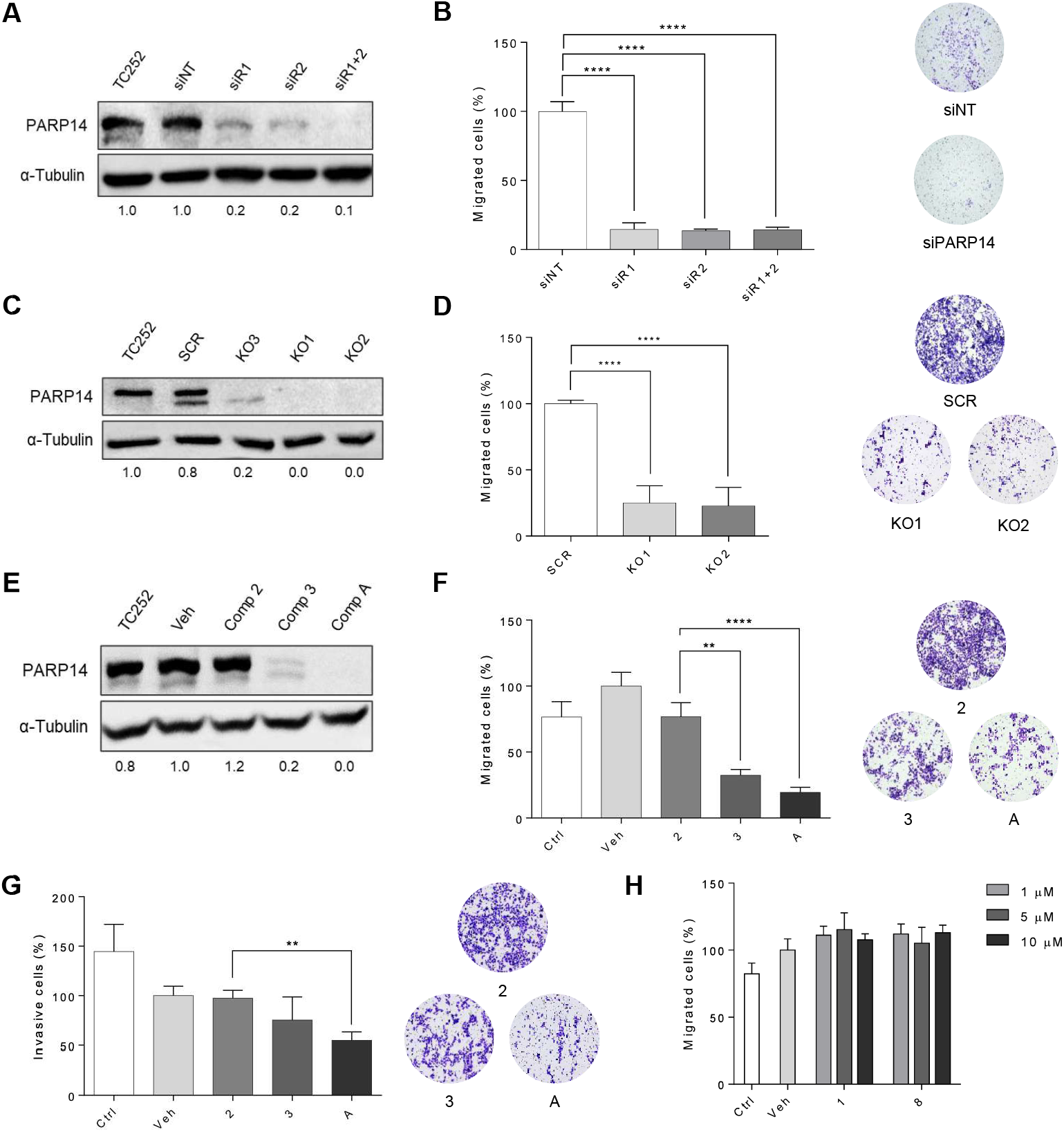
PARP14 depletion impairs EwS migration and invasion in vitro. **A)** PARP14 knock-down protein levels measured by WB 48 h post-transfection in TC252 and cell line. α-Tubulin was used as loading control. Protein levels are relative to cells treated with a non-targeting control siRNA (siNT). Two different siRNAs were tested (siR1 and siR2) as well as the combination of both (siR1+2). **B)** Migration transwell assay in TC252 metastatic EwS cells after PARP14 siRNA-mediated transient knock-down. Pictures of the lower part of the membranes and their quantification. 48 h of migration. **C)** PARP14 CRISPR-Cas9 mediated knock-out model protein levels by WB in TC252 and cell line. α-Tubulin was used as loading control. Protein levels of CRISPR-Cas9 clones are relative to TC252. **D)** Migration transwell assay in TC252 PARP14 CRISPR Cas9-mediated KO clones. Pictures of the lower part of the membranes and their quantification. 48 h of migration. **E)** PARP14 protein levels measured by WB 48 h after 10 nM PROTACs treatment [Compound 3 (RBN012811), Compound A and inactive Compound 2 (RBN013527)] in TC252 cell line. α-Tubulin was used as loading control. Protein levels of PROTAC treatments are relative to TC252 treated with vehicle (DMSO). **F)** Migration and **G)** Invasion transwell assay in metastatic TC252 cells after PARP14 PROTACs-mediated degradation. Pictures of the lower part of the membranes and their quantification. 48 h of migration/invasion. **H)** Migration transwell assay in TC252 metastatic cells after PARP14 catalytic inhibition using different concentrations of Compound 8 (RBN012759) and inactive Compound 1. Statistical analyses were made using one-way ANOVA test. ^**^p<0.01; ^****^p<0.00 1.

To further validate these findings, EwS cells migration and invasion abilities were assessed following PARP14 depletion using proteolysis-targeting chimeras (PROTACs). PARP14 degraders Compound 3 (RBN012811) and its optimized analog Compound A were used, which bind to PARP14’s NAO-binding site and recruit cereblon to ubiquitinate it and selectively target it for degradation via the proteasome system. Treatment of metastatic TC252 and MHHES1 cells with Compounds 3 or A depleted PARP14 protein levels at concentrations as low as 10 nM (Fig. 2E; Fig. S2E), resulting in a marked reduction in both migration and invasion compared with DMSO-treated cells (Veh) and those treated with the inactive analog Compound 2 (RBN013527) (Fig. 2F-G; Fig. S2F).

To determine whether the pro-metastatic role of PARP14 was dependent on its enzymatic activity, a selective catalytic inhibitor (Compound 8; RBN012759) was tested versus its inactive form (Compound 1). Notably, inhibition of PARP14 catalytic activity did not affect the migratory ability of TC252 cells across a range of concentrations (Fig. 2H), suggesting that PARP14 promotes EwS metastasis through a mechanism independent of its enzymatic function.

To evaluate whether PARP14 depletion impacts other tumorigenic phenotypes, we examined cell proliferation and clonogenic potential following PARP14 loss-of-function. Neither PARP14 degradation by PROTACs nor inhibition of its catalytic activity with Compound 8 affected TC252 cell proliferation or colony-forming ability (Fig.S3A-D).

### Genetic and pharmacologic PARP14 inhibition influences EwS metastasis *in vivo*

To determine whether the observed effects on migration and invasion *in vitro* translate into altered tumor growth and metastatic behavior *in vivo*, we next evaluated TC252 PARP14 KO clones (K01 and K02) in our spontaneous metastasis xenograft model. Loss of PARP14 was associated with delayed tumor growth, as indicated by the increased interval between primary tumor resection surgeries in mice injected with KO clones compared with those injected with parental TC252 cells (Fig. 3A). Further, mice bearing tumors derived from the K01 clone exhibited significantly fewer *ex vivo* lung metastatic lesions (Fig. 3B). Notably, IHC staining of lung metastases revealed PARP14 re-expression in lesions from KO-bearing mice (Fig. S4A).

**Figure 3.**
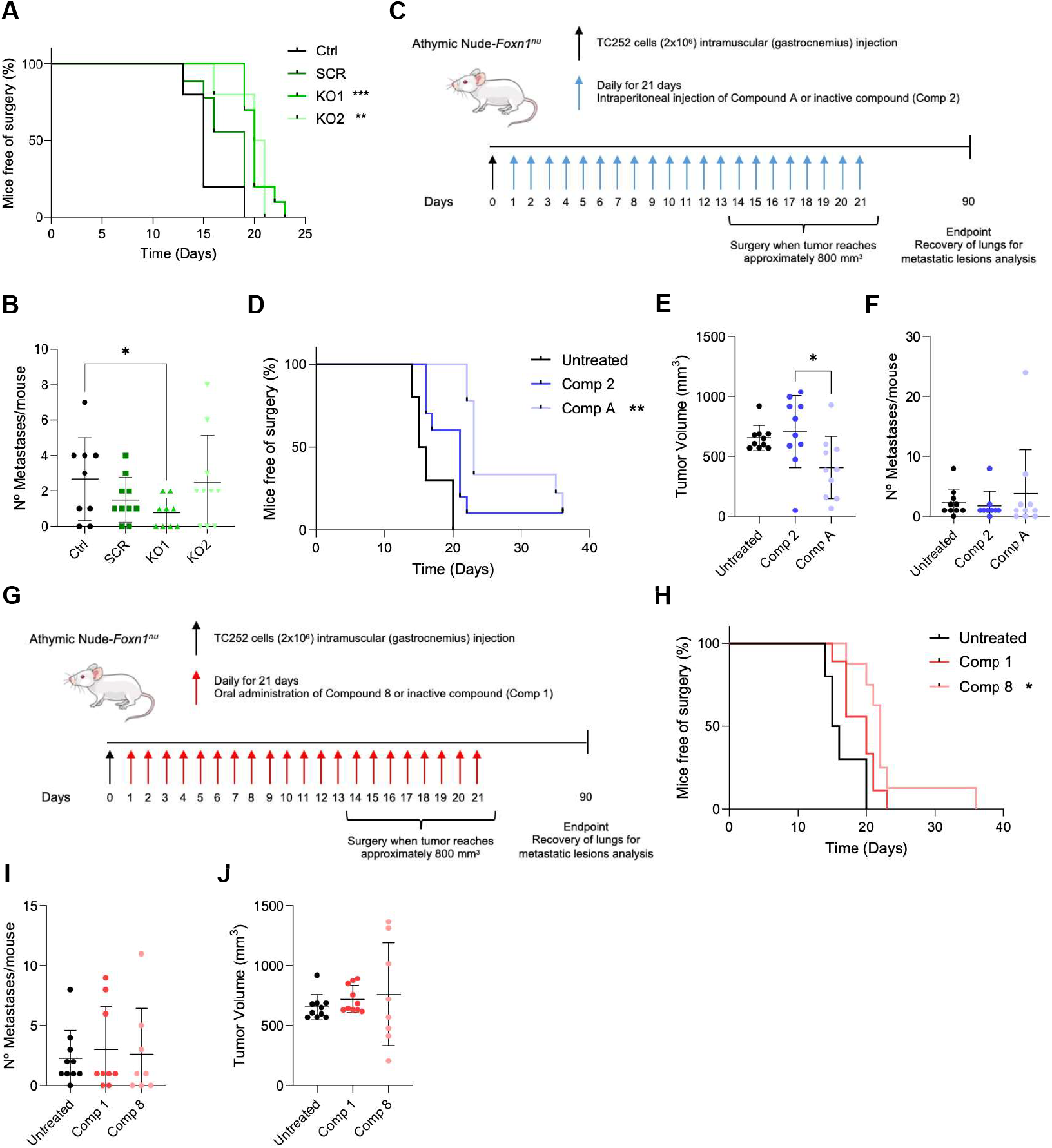
PARP14 depletion delays EwS tumorigenesis and partly reduces metastasis in vivo. **A)** Kaplan-Meier curves depicting mice-free of surgery upon time (days) of mice injected with TC252 control cells, CRISPR-Cas9 mediated scrambled (SCR) or CRISPR-Cas9 mediated clones (KO1 and KO2), n=10 mice for each condition and **B)** Total lung metastases obtained per mouse considering both macro and micro metastases. **C)** Diagram depicting the main steps of the spontaneous metastasis mouse model treated with PARP14 PROTAC Compound A. TC252 cells (2’106) were injected in the gastrocnemius muscle of athymic mice. Compound A or its inactive version (Comp 2) were daily injected (IP) until day 21. Primary tumors were collected when reached 800 mm3. At day 90 (endpoint), mice were euthanized and lung metastases collected. **D)** Kaplan-Meier curves depicting mice-free of surgery upon time (days) of mice injected with TC252 cells and treated daily for 21 days with PROTAC Compound A or its inactive form (Comp 2). **E)** Tumor volumes of the different conditions at day 21 of treatment (last day of treatment administration). **F)** Total lung metastases obtained per mouse considering both macro and micro metastases. **G)** Diagram depicting the main steps of the spontaneous metastasis mouse model treated with PARP14 catalytic inhibitor Compound 8. TC252 cells (2’106) were injected in the gastrocnemius muscle of athymic mice. Compound 8 or its inactive version (Comp 1) were daily administered (OA) until day 21. Primary tumors were collected when reached 800 mm3. At day 90 (endpoint), mice were euthanized and lung metastases collected. **H)** Kaplan-Meier curves depicting mice-free of surgery upon time (days) of mice injected with TC252 cells and treated daily for 21 days with catalytic inhibitor Compound 8 or its inactive form (Comp 1). **I)** Total lung metastases obtained per mouse considering both macro and micro metastases. **J)** Tumor volumes of the different conditions at day 21 of treatment (last day of treatment administration). Statistical analyses were made using Mantel-Cox and one-way ANOVA test. ^*^p<0.05; ^**^p<0.01; ^***^p<0.001.

PARP14 PROTACs were also tested *in vivo* using the same mouse model. TC252 cells were injected into the gastrocnemius muscle of mice and treated daily during 21 days by subcutaneous administration of either Compound A or its inactive analog (Compound 2; Fig. 3C). Consistent with the genetic KO results, Compound A-treated mice displayed a significant delay in tumor growth compared with those treated with the inactive compound (Fig. 3D). Interestingly, tumors from Compound A-treated mice were significantly smaller than those from both control and Vehicle 2 groups at day 21 (last day of drug administration) (Fig. 3E). Additionally, a lower tendency of developing lung metastases was observed in mice treated with Compound A (except from an outlier); however, this difference did not reach statistical significance likely due to early termination of treatment (Fig. 3F).

The catalytic inhibitor was also assessed *in vivo*. Here, mice injected with TC252 cells were treated daily for 21 days by oral administration of either Compound 8 or its inactive analog (Compound 1; Fig. 3G). A modest reduction in tumor growth was observed in Compound 8-treated mice (Fig. 3H); however, no significant differences in lung metastasis formation were detected between the two treatment arms (Fig. 3I). Similarly, tumor volumes at day 21 did not differ significantly among the three groups (Fig. 3J).

To determine whether PARP14 expression was restored following treatment cessation, we obtained primary cultures from lung metastases of the different treatment groups (Fig. S4B). Interestingly, all recovered metastases displayed notably higher PARP14 expression compared with the parental TC252 cells, suggesting re-expression upon drug withdrawal (Fig. S4C).

### PARP14 upregulation increases migration and invasion in EwS

As previously reported, IFNγ can induce PARP14 expression in multiple cell types. Consistent with this, treatment with IFNγ led to dose-dependent upregulation of PARP14 in EwS cells with (TC252) or without (A673) basal expression (Fig. 4A-B). This increase in PARP14 levels was accompanied by enhanced migration of TC252 cells, which was at least partially dependent on PARP14, as IFNγ-treated KO cells failed to recover migration levels of SCR control cells (Fig. 4C).

**Figure 4.**
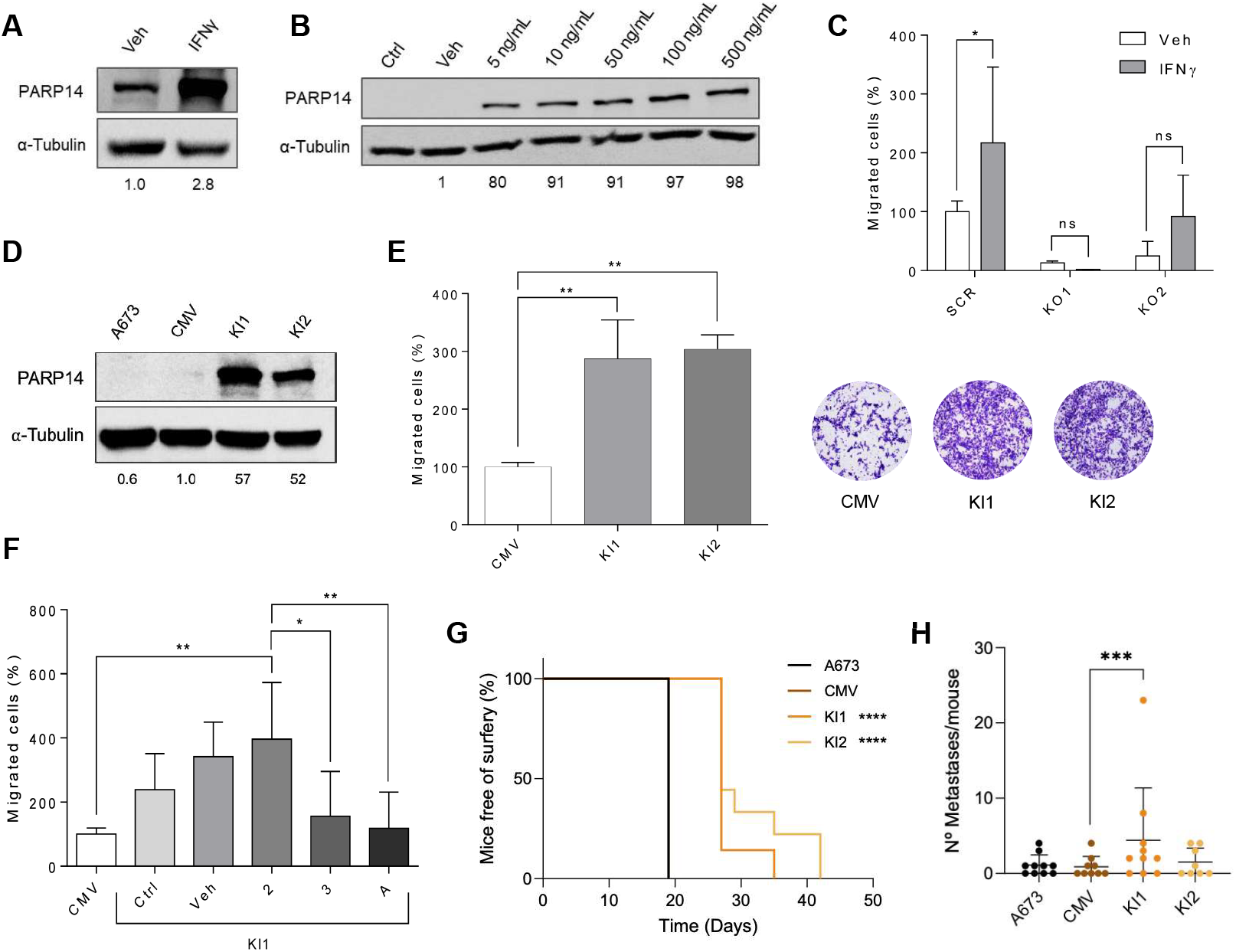
PARP14 upregulation increases migration and invasion in EwS. **A)** PARP14 protein levels measured by WB after 16 h of treatment with 100 ng/ml of IFNγ in TC252 cell line. α-Tubulin was used as loading control. PARP14 protein levels are relative to TC252 treated with vehicle (Veh). **B)** PARP14 protein levels measured by WB after 16 h of treatment with different concentrations of IFNγ in A673 cell line. α-Tubulin was used as loading control. PARP14 protein levels are relative to A673 treated with vehicle (Veh). **C)** Migration transwell assay of TC252 PARP14 KO stable model after a 16 h IFNγ 100 ng/ml pre-treatment. Picture of the lower part of the membranes and their quantification. Migration levels are relative to TC252 SCR treated with vehicle (Veh). 48 h of migration. **D)** PARP14 Kl model protein levels by WB in A673 and cell line. α-Tubulin was used as loading control. Protein levels of Kl clones are relative to A673 transfected with a control vector (CMV). **E)** Migration transwell assay of A673 PARP14 Kl stable model. Picture of the lower part of the membranes and their quantification. Clones KI1 and KI2 migration levels are relative to A673 transfected with a CMV control vector. 24 h of migration. **F)** Migration transwell assay of A673 PARP14 Kl1 clone after a 24 h PROTACs 10 nM [Compound 3 (RBN012811), Compound A and inactive Compound 2 (RBN013527)] pre-treatment. Picture of the lower part of the membranes and their quantification. Migration levels are relative to A673 transfected with a CMV control vector. 24 h of migration. **G)** Kaplan-Meier curves depicting mice-free of surgery upon time (days) of mice injected with A673 control cells, A673 transfected with a CMV control vector (CMV) or PARP14 Kl stable clones (Kl1 and Kl2), n=10 mice for each condition. **H)** Total lung metastases obtained per mouse considering both macro and micro metastases. Statistical analyses were made using unpaired t-test with Welch’s correction, Mantel-Cox and one-way ANOVA test. ^*^p<0.05; ^**^p<0.01.

To further validate the phenotypic effects of PARP14 upregulation, a stable FLAG-tagged PARP14 knock-in (Kl) model was developed in A673 cell line (Fig. 4D). Consistent with our loss-of-function findings, PARP14 overexpression significantly enhanced cell migration (Fig. 4E). Importantly, this effect was PARP14-dependent, as treatment of Kl 1 clone with PARP14 PROTACs restored migration levels to those of the control groups (Fig. 4F). In line with the KO model, PARP14 overexpression did not affect colony formation or proliferation abilities of A673 cells (Fig. SA-8).

Finally, the PARP14 Kl model was also assessed *in vivo*. Mice injected with cells overexpressing PARP14 exhibited a modest delay in tumor growth (Fig. 4G). Further, mice bearing tumors derived from the Kl1 clone exhibited significantly higher *ex vivo* lung metastatic lesions (Fig. 4H), in line with the functional effects observed *in vitro*.

### PARP14 plays a role in tumor inflammation and the epigenomic landscape of EwS

To investigate the mechanisms by which PARP14 promotes metastasis in EwS, RNA-seq analysis was performed comparing TC252 KO clones (KO1 and KO2) versus control cells (TC252 and scrambled (SCR)). Heatmap visualization revealed distinct transcriptional landscapes between these 2 groups of samples (Fig. 5A; Supplementary Table 1). Differential expression analysis identified 606 upregulated and 471 downregulated transcripts in the KO clones compared with controls (LFC > |1|, adjusted p < 0.0001; Fig. 5B).

**Figure 5.**
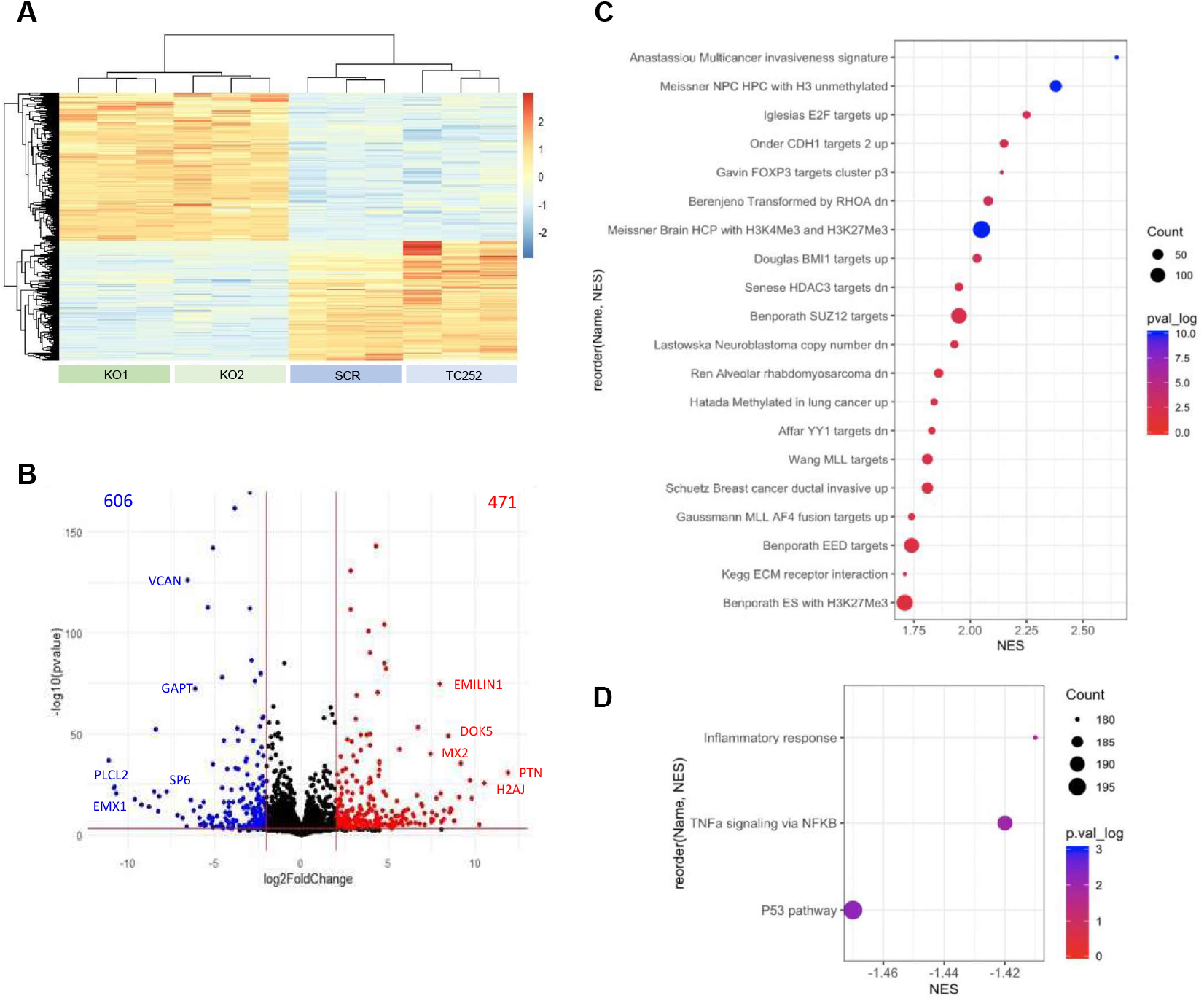
PARP14 plays a role in tumor inflammation and the epigenomic landscape of EwS. **A)** Transcriptomic profiling of TC252 control and scrambled (SCR) cells versus CRISPR-Cas9 mediated knock-out clones G2_11 and G2_24 revealed differential expression profile between these 2 groups of samples. **B)** Volcano plot of transcriptomic results showing 606 transcripts enriched in knock-out samples (blue) compared to 471 transcripts enriched in control cells (red) (LFC > | 1 |, adjusted p < 0.0001). **C)** GSEA pre-ranked analysis showing gene sets downregulated in PARP14 knock-out samples when compared to controls using C2 datasets. **D)** GSEA pre-ranked analysis showing gene sets enriched in PARP14 knock-out samples when compared to controls using Hallmarks datasets.

Pre-ranked Gene Set Enrichment Analysis (GSEA) using C2 curated gene sets revealed an invasive signature as the most significantly downregulated in PARP14 KO cells (Fig. 5C). Also, a high number of datasets related with methylation marks (i.e. Meissner NPC HPC with H3 unmethylated, Meissner Brain HCP with H3K4Me3 and H3K24Me3, Sense HDAC3 targets down) and chromatin remodeling complexes such as the polycomb group complex (i.e. Douglas BMl1 targets up, Benporath SUZ12 targets, Benporath EEO targets) were overlapping with PARP14 depletion, suggesting that PARP14 may influence epigenomic regulation in EwS. Additionally, pre-ranked GSEA using Hallmarks gene sets revealed p53 signaling, TNFa signaling via NF-KB and inflammatory response as pathways upregulated upon PARP14 depletion (Fig. 5D).

### PARP14 acts as a chromatin associated co-regulator in the nucleus of EwS cells

Previous studies have reported PARP14 localization in both cytoplasmic and nuclear compartments. Consistently, subcellular fractionation of A673 cells treated with IFNγ or stably overexpressing PARP14 revealed its distribution across the cytoplasmic, nuclear, and chromatin-bound fractions, each accounting for approximately one-third of total PARP14 (Fig. 6A). In addition, since the A673 PARP14 Kl model includes a BFP2 reporter, confocal microscopy confirmed PARP14 localization in both cytoplasmic and nuclear compartments (Fig. S5C). Notably, analysis of PARP14 staining in EwS patient samples (Fig. 1E) showed a correlation of metastasis presence and PARP14 location in the nucleus (data not shown), suggesting that its pro-metastatic role may extend beyond its canonical function in the cytoplasm.

**Figure 6.**
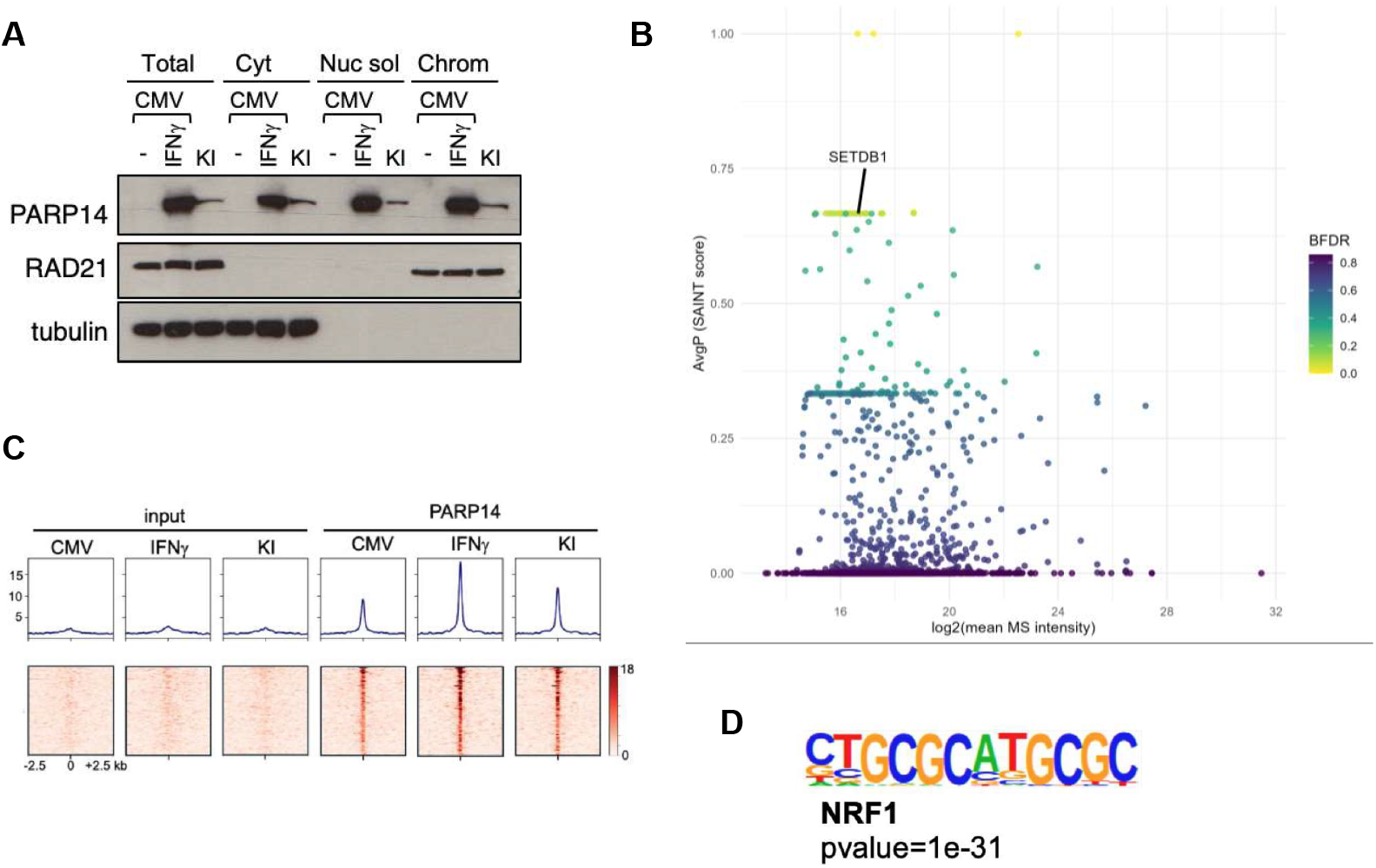
PARP14 acts as a chromatin associated co-regulator in the nucleus of EwS cells. **A)** PARP14 protein levels after subcellular fractionation in control A673 CMV, IFNγ treated-A673 CMV and A673 PARP14 Kl1 cells. RAD21 and a-Tubulin used as controls of chromatin and cytoplasm fractions, respectively. Total: whole cell extract. Cyt: cytoplasmic fraction. Nuc sol: nuclear soluble fraction. Chrom: chromatin fraction. **B)** PARP14 nuclear interactors identified by pull-down/MS. Candidate interactor intensity on the x-axis and software-derived score (SAINT) on the y-axis. Color indicates the Bayesian false discovery rate (BFDR). **C)** Read density (top) and heatmaps (bottom) around 79 peaks identified for PARP14 binding. Inputs are also shown. **D)** Top motif identified in the binding sites of PARP14.

To explore how nuclear PARP14 might contribute to the epigenomic remodeling observed in EwS cells, pull-down assays followed by mass spectrometry on nuclear and whole extracts from control A673 (CMV) and PARP14 Kl1 cells were performed. This analysis identified a direct interaction between PARP14 and chromatin regulators such as SETDB1, NSD1, HDAC8, NCAPG2 and RCOR3 (Fig. 6B; Supplementary Table 2). Other PARP14-binding partners included JCAD, SLC26A2, CD2AP, THBS3, and DNMBP, all linked to cell-cell junctions, cytoskeletal signaling, and extracellular matrix remodeling in other contexts.

In an attempt to deep into the nuclear mechanisms by which PARP14 might modulate its aggressiveness, we performed a ChlP-seq experiment in A673 CMV and PARP14 Kl cells. A673 cells treated with IFNγ were also included as an additional control. As expected for a transcriptional regulator that does not bind directly to DNA, very few *bona fide* PARP14 peaks (n=79) were identified. The strength of the peaks correlated with the levels of PARP14 present in the three conditions (Fig. 6C) and most of them (>85%) were located at gene promoters or in adjacent CpG islands (Supplementary Table 3). Motif-based sequence analysis revealed a highly significant enrichment in the binding site of the transcriptional factor NRF1, which was present in more than 72% of the PARP14 binding sites (Fig. 6D). NRF1 regulates genes critical for mitochondrial biogenesis, metabolism and cellular growth. This result suggests that PARP14 could modulate NRF1 transcriptional activity and participate in the metabolic shift that characterizes the more aggressive phenotype of metastatic EwS. To our knowledge, this is the first study describing PARP14 acting as a transcriptional co-regulator in this disease.

Together, our results uncover a nuclear function of PARP14 as a chromatin-associated co-regulator and suggest that it may influence transcriptional programs relevant to EwS progression. Combined with its interactions with proteins involved in cytoskeletal organization and cell-cell junctions, these findings point to a multifaceted role for PARP14, potentially acting through both nuclear and cytoplasmic pathways to promote metastatic behavior. This highlights PARP14 as a versatile regulator in EwS and provides a strong rationale for further investigation into the mechanisms by which it contributes to disease progression.

## DISCUSSION

Despite all the efforts of multidisciplinary teams trying to untangle the molecular and biological mechanisms behind EwS progression, the successful treatment of young patients affected by this disease remains a major challenge. This is particularly true for metastatic patients, who still lack specific therapeutic options for their condition^2^.

Trying to shed some light on genes and pathways involved in EwS metastasis, we started this project comparing primary tumors and lung metastases from our spontaneous metastasis xenograft mouse model^4, 12^ identifying PARP14 as significantly and consistently upregulated in metastatic samples. PARP14 transient and stable loss-of-function significantly impaired migration ability of EwS cells in Boyden chamber assays, suggesting a potential role of this protein in EwS progression. Although PARP14 did not appear to be involved in regulating cell proliferation or tumorigenesis *in vitro*, we observed a pronounced delay in tumor growth *in vivo* upon PARP14 depletion, either through CRISPR-Cas9 knockout or PROTAC-mediated degradation. These results suggest that, in a physiological context that more accurately recapitulates tumor features, PARP14 contributes to EwS tumor growth.

Regarding the metastatic process, mice injected with PARP14 KO clones or treated with PROTACs still developed metastases. This was most likely due to PARP14 re-expression observed in these metastatic lesions, as suggested by immunoblot analysis of metastasis-derived primary cell cultures or in situ IHC staining of lung tissues.

Together with the *in vitro* findings, these results support our hypothesis that PARP14 promotes EwS cell dissemination. The TME and the complex interactions between extracellular components, stromal elements, and malignant cells, can profoundly influence tumor behavior and adaptation. Notably, PARP14 has been associated with inflammation and TME modulation^10,24-26^ and is highly expressed in pro-tumor macrophages and B cells^8– 10^. Therefore, its study in isolated *in vitro* systems may only partially capture its role *in vivo*, where immune and stromal players exert major influences.

Interestingly, PARP14-mediated EwS progression appeared to be, in our model, independent of its catalytic function (ADP-ribosylation of target proteins), as inhibition of its enzymatic function did not affect any of the phenotypic properties analyzed *in vitro* or *in vivo*. Of note, PARP14 is known to conduct its enzymatic function primarily in the cytoplasm, as it performs a post-translational modification of its targets^5^. However, we have also detected its presence in the nuclear soluble and chromatin fractions, consistent with previous reports^27-29^. To definitively confirm the catalytic-independent function of PARP14 in EwS, site-directed mutagenesis of the catalytic domain will be required.

PARP14 expression has been reported to be induced by both type I and II interferons^10,11,28^. We observed a strong and consistent upregulation of PARP14 mRNA and protein levels following IFNγ treatment, both in cells with and without basal expression levels. This IFNγ-driven PARP14 induction was followed by a significant increase in EwS cells migration. We therefore hypothesize that IFNγ-driven PARP14 upregulation in a subset of primary tumor cells could facilitate their escape from the primary site, circulation, extravasation, and colonization of distant organs, ultimately leading to macrometastatic lesions. In line with this, a growing body of evidence implicates IFNγ in promoting tumorigenesis, angiogenesis and immune evasion, thereby contributing to disease progression^30-33^. Macrophages may be the source of IFNγ production in our model^34-36^. Indeed, immunosuppressive M2-like type macrophages are the most abundant immune infiltrates in EwS tumors, and their presence correlates with poor prognosis^37-39^. Interestingly, pharmacological inhibition of macrophages using CNl-1493 effectively reduced EwS metastasis in an *in vivo* mouse model through impairment of cell extravasation^40^, further highlighting the pro-tumoral role of macrophages in this disease. More research is needed to uncover this potential macrophage-lFNγ-PARP14 axis in EwS and its possible targeting as an approach to overcome or even avoid metastasis in these tumors.

In the last part of our study, we focused our attention on understanding how PARP14 exerts its pro-metastatic function in EwS. RNA-seq analysis comparing parental TC252 or SCR cells with two independent PARP14 KO clones revealed upregulation of TNFα signaling via NF-κB and inflammatory response pathways upon PARP14 loss, further reinforcing the link between PARP14, inflammation, and TME modulation. Additionally, several datasets related to methylation marks and chromatin remodeling complexes such as the polycomb group complex were enriched among genes downregulated upon PARP14 depletion. These findings suggest a potential role of PARP14 in shaping the epigenetic landscape of EwS cells during disease progression.

Mechanistically, we identified a direct interaction between PARP14 and SETDB1, a histone methyltransferase associated with transcriptional repression and previously reported to be a mediator of immune escape^41,42^. Complementary ChIP analyses revealed PARP14 enrichment at promoters of NRF1-regulated genes. This transcription factor has been showed to be involved in metabolism, mitochondrial biogenesis and cellular growth^43,44^. To our knowledge, this is the first study to attribute to PARP14 a transcriptional co-regulatory role in EwS, suggesting that it may influence gene networks relevant to disease progression through chromatin-associated functions. In addition to SETDB1, our interactome analysis identified other PARP14-binding partners, including JCAD, CD2AP, THBS3 and DNMBP, all related to cell-cell junctions, cytoskeletal signaling and ECM remodeling^45– 48^. These findings underscore the possibility that PARP14 operates at multiple cellular levels and compartments to coordinate pro-metastatic programs. Indeed, PARP14 has previously been reported as a component of focal adhesion complexes required for proper cell motility and adhesion dynamics^49^, further supporting its multifaceted contribution to EwS dissemination.

In conclusion, our study sheds light on the role of PARP14 in promoting EwS progression. By integrating *in vitro, in vivo*, transcriptomic and epigenetic data, we propose that PARP14 supports EwS tumor growth and dissemination, potentially through mechanisms beyond its catalytic activity, involving TME modulation and epigenetic regulation. Moreover, we provide the first evidence that PARP14 can act as a transcriptional co-regulator, directly engaging with chromatin at promoters enriched for NRF1 motifs and influencing NRF1-dependent gene expression.

Given its consistent association with aggressive disease and its potential value as a biomarker, PARP14 may represent a promising candidate for both prognostic assessment and therapeutic intervention in EwS. We hope our findings contribute to a better understanding of the molecular drivers of EwS metastasis and ultimately aid in the development of more effective treatment strategies for patients.

## ACKNOWLEDGEMENTS

This work was supported by grants from Institute de Salud Carlos Ill and FEDER (CES12/021). Ministerio de Ciencia, lnnovación y Universidades (RTl2018-094787-B-I00; PID2021-122828OB-I00). AGAUR (2017SGR332). La Marató de TV3 (318/C/2019). CERCA Programme/Generalitat de Catalunya. Asociación Pablo Ugarte. MCIN/AEl/10.13039/501100011033 and ERDF “A way of making Europe” grant PID2022-139333NB-100 (to A.L.).

## AUTHOR CONTRIBUTIONS

Conceptualization: OMT; methodology: SSS, MCB, JJBS, RSP, AC, AL and RLA; investigation and interpretation: SSS, MCB, NMV, EC, IJS, JJBS, SGL, MGL, PM, LSV, MN, RSP, AC, AL, RLA and OMT; resources: SML, JM, MN; writing - original draft preparation, SSS; writing - review and editing: SSS, AC, AL, RLA and OMT; visualization: SSS, EC, MCB, JJBS, AC and AL; funding acquisition; AC, AL and OMT; software: MCB, EC, IJS, JJBS, DGL and RSP.

## DATA AND CODE ACCESSIBILITY

The mass spectrometry proteomics data has been deposited to the ProteomeXchange Consortium via the PRIDE^50^ partner repository with the dataset identifier PXD072018. Log in details can be provided upon reviewer request.

