## Supplemental figures for "Integrative multi-omics reveals PARP14 as a key IFNγ-regulated mediator of metastatic progression in Ewing sarcoma"

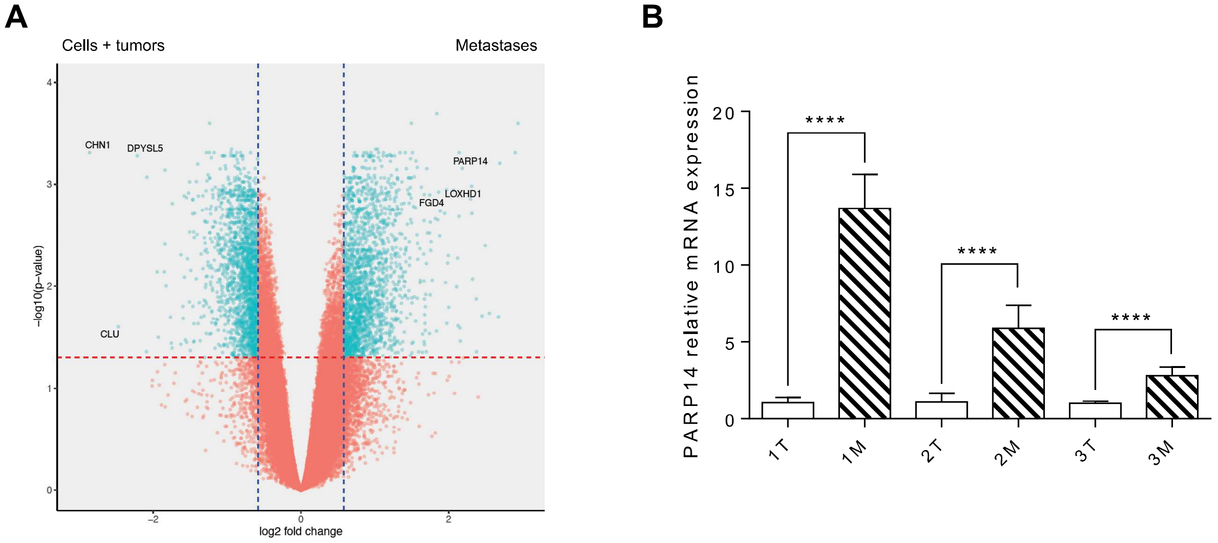


Supplementary Figure 1

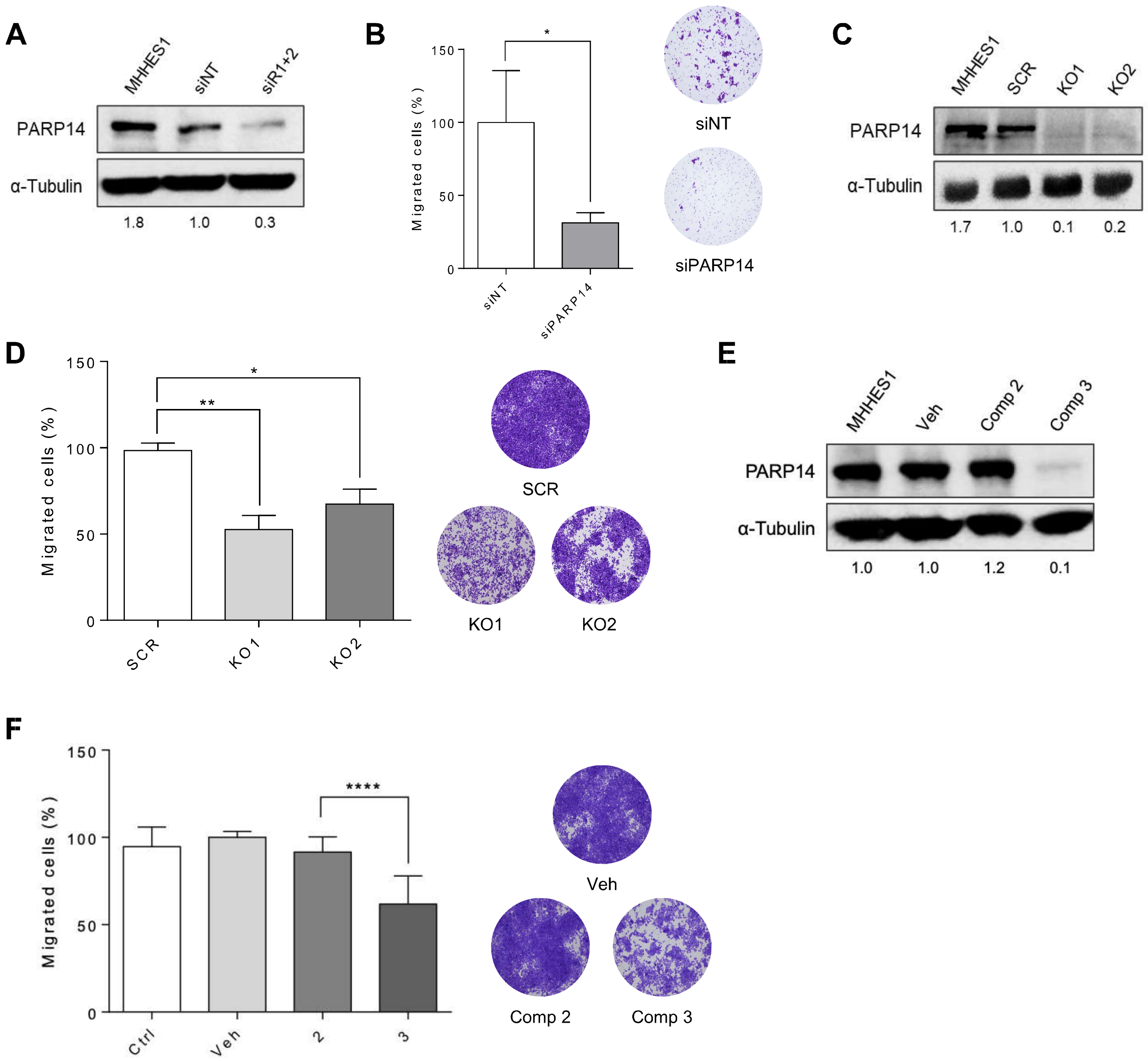


Supplementary Figure 2

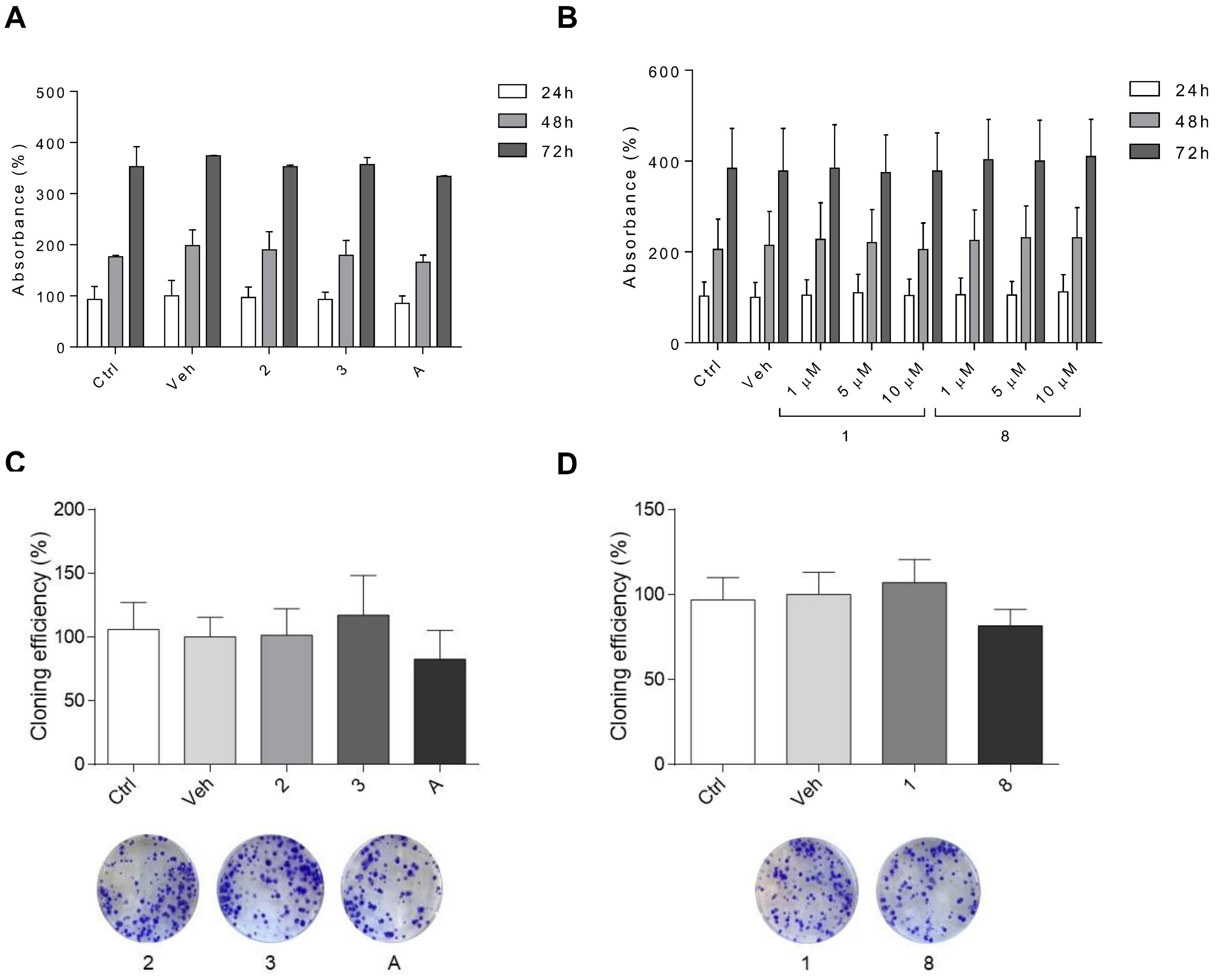


Supplementary Figure 3

**A**

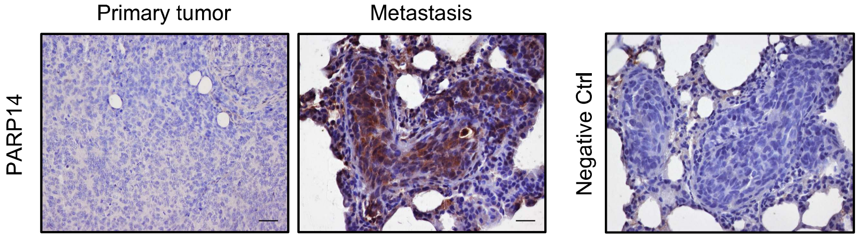


**B**

| Primary culture | Tumor subtype | Cell line of origin | Treatment |
|---|---|---|---|
| **Met2TC** | EwS | TC252 | Untreated |
| **Met3TC** | EwS | TC252 | Compound 8 |
| **Met4TC** | EwS | TC252 | Compound 1 |
| **Met5TC** | EwS | TC252 | Compound A |

**C**

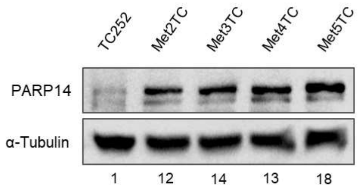


Supplementary Figure 4

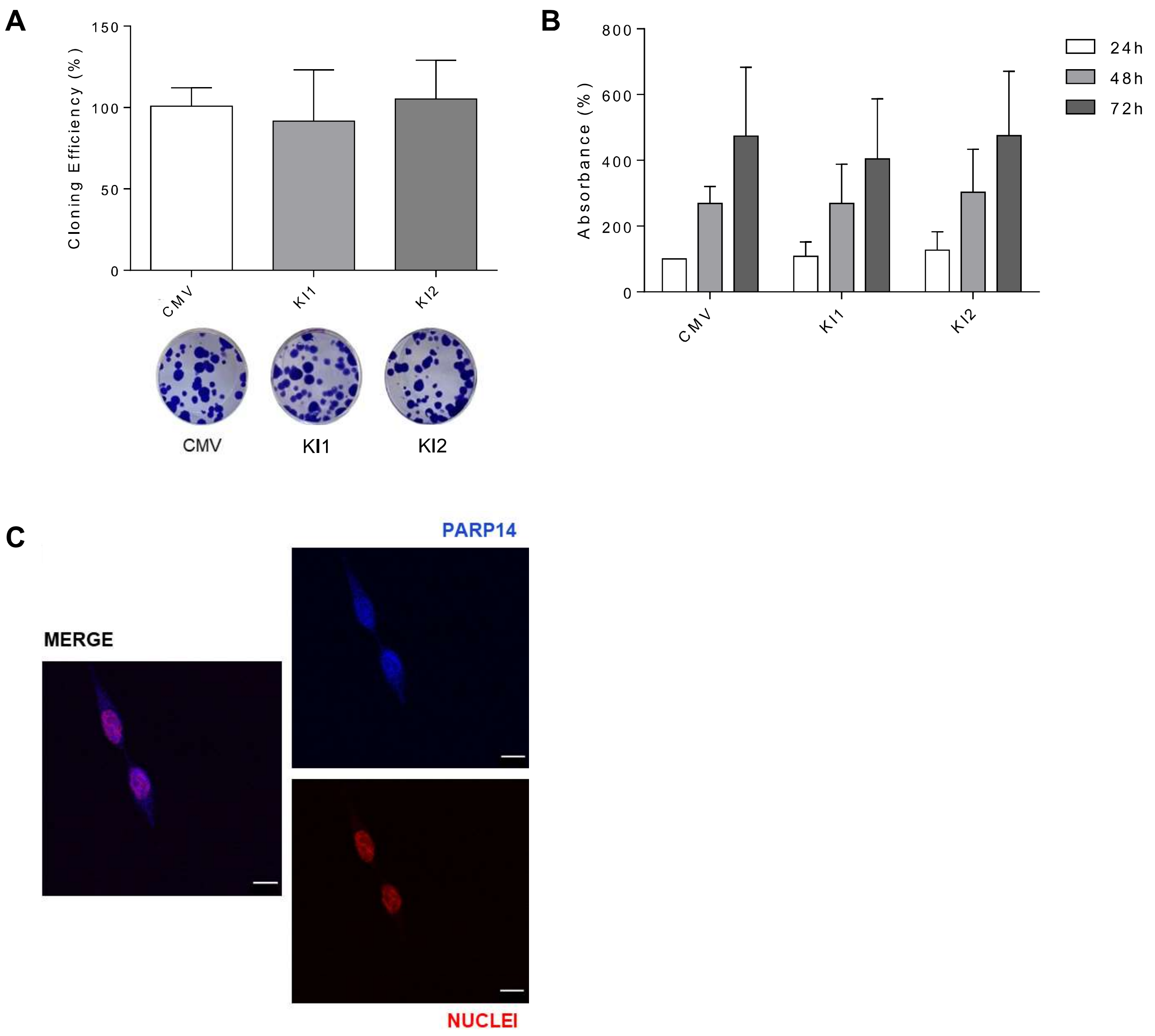


Supplementary Figure 5
