## Supplemental figure legends for "Integrative multi-omics reveals PARP14 as a key IFNγ-regulated mediator of metastatic progression in Ewing sarcoma"

### SUPPLEMENTARY FIGURE LEGENDS

**Supplementary Figure 1. A)** Volcano plot of differential expression analyses comparing A673 primary tumors and cells (n=4+4) versus metastases (n=12). In blue, all transcripts that are significant (LFC >|0.58|, adjusted p-value <0.05). 2075 transcripts enriched in metastases samples compared to 2062 transcripts enriched in Cells+Tumors. PARP14 and other examples of DEGs are shown. **B)** PARP14 mRNA relative levels of TC252-derived xenograft samples by RT-qPCR. PARP14 mRNA levels of each metastasis are relative to their corresponding tumor levels (normalized to 1). T: tumor; M: metastasis. Statistical analyses were made using unpaired t-test with Welch’s correction. ****p<0.0001.

**Supplementary Figure 2. A)** PARP14 knock-down protein levels measured by WB 48 h post-transfection in MHHES1 cell line. α-Tubulin was used as loading control. Protein levels are relative to cells treated with a non-targeting control siRNA (siNT). Two different siRNAs were tested in combination (siR1 and siR2). **B)** Migration transwell assay in MHHES1 metastatic EwS cells after PARP14 siRNA-mediated transient knock-down. Picture of the lower part of the membranes and their quantification. 48 h of migration. **C)** PARP14 CRISPR-Cas9 mediated KO model protein levels by WB in MHHES1 cell line. α-Tubulin was used as loading control. Protein levels of CRISPR-Cas9 clones are relative to MHHES1 transfected with a scrambled vector (SCR). **D)** Migration transwell assay in MHHES1 metastatic EwS cells after PARP14 CRISPR-Cas9-mediated stable KO. Picture of the lower part of the membranes and their quantification. 48 h of migration. **E)** PARP14 protein levels measured by WB 48 h after 10 nM PROTACs treatment in MHHES1 cell line. α-Tubulin was used as loading control. Protein levels of PROTAC treatments are relative to MHHES1 treated with vehicle (DMSO). **F)** Migration transwell assay in MHHES1 metastatic EwS cells after PARP14 PROTACs-mediated degradation [Compound 3 (RBN012811) and inactive Compound 2 (RBN013527)]. Pictures of the lower part of the membranes and their quantification. 48 h of migration. Statistical analyses were made using unpaired t-test with Welch’s correction and one-way ANOVA test. *p<0.05; **p<0.01; ****p<0.0001.

**Supplementary Figure 3. A)** Proliferation of TC252 cells treated with 10 nM of PARP14 PROTACs [Compound 3 (RBN012811), Compound A and inactive Compound 2 (RBN013527)] or **B)** different concentrations of PARP14 inhibitors [Compound 8 (RBN012759) and inactive Compound 1] measured by MTT after 24, 48 and 72 h of treatment. Values are relative to TC252 cells treated with vehicle (Veh) after 24 h of treatment. **C)** Colony formation assay in TC252 cells treated with 10 nM of PARP14

PROTACs or **D)** 1 μM of PARP14 inhibitors 24 h after cells seeding. Pictures of the wells (14 days after seeding) and their quantification. Values are relative to TC252 cells treated with vehicle (Veh). Statistical analyses were made using one-way ANOVA test.

**Supplementary Figure 4. A)** PARP14 protein expression assessed by immunohistochemistry staining of primary tumor and lung metastasis from xenograft mouse model bearing a tumor derived from TC252 PARP14 KO2 clone cells. Scale bar = 50 μm. **B)** Metastasis-derived cell lines obtained directly from mouse metastatic lesions as a primary culture. Tumor type, cell line from which it is derived and treatment administered to each mouse (if applies) are described. **C)** PARP14 protein relative levels on xenograft-derived metastatic cell lines by WB. α-Tubulin was used as loading control. PARP14 protein levels of each metastatic cell line are relative to TC252.

**Supplementary Figure 5. A)** Colony formation assay in A673 PARP14 KI model. Pictures of the wells (14 days after seeding) and their quantification. Values are relative to A673 transfected with a CMV control vector. **E)** Proliferation of A673 PARP14 KI model measured by MTT after 24, 48 and 72 h of seeding. Values are relative to A673 transfected with a CMV control vector after 24 h of seeding. **C)** Representative images showing PARP14 (blue) by confocal microscopy on a knock-in model in A673 cells. Nuclei in red. Scale bar: 10 μm. Statistical analysis were made using one-way ANOVA test.
